# High-Efficiency Capture and Proteomic Analysis of Plasma-Derived Extracellular Vesicles through Affinity Purification

**DOI:** 10.1101/2024.08.01.605729

**Authors:** Gui-Yuan Zhang, Cheng-Xiao Ma, Le Ma, Dong Wei, Ya-Nan Wu, Ying Li, Zhe-Hui Xu, Yu-Feng Liu, Yu-Han Cai, Evan Yi-Wen Yu, Ye-Fei Zhu, Hao Zhang

## Abstract

Plasma-derived extracellular vesicles (EVs) are promising sources of biomarkers. It is still a challenge to isolate EVs from a small amount of human plasma for downstream proteomic analysis. The separation process is hindered by contamination with high-abundance blood proteins and lipoprotein particles, which adversely impact proteomic analyses. Moreover, although EVs immune-separation via magnetic beads often integrates with flow sorting and western blotting (WB), it lacks compatibility with nanoparticle tracking analysis (NTA) and proteomic analysis. To address these issues, we have developed a functional affinity magnetic bead, EVlent (**E**xtracellular **V**esicles iso**L**ated **E**fficiently, **N**aturally, and **T**otally), enabling the rapid and efficient separation of EVs from plasma. By optimizing the quantities of magnetic beads and plasma used, we characterized the isolated EVs through WB, NTA, and transmission electron microscopy (TEM), showing a successfully separation of EVs from plasma. Proteomic analysis of these EVs identified over 2,000 proteins and 15,000 peptides from just 100 μL of plasma, and nearly 1,000 proteins from trace samples as small as 5 μL. Additionally, this isolation method significantly reduced contaminants, including plasma proteins and lipoproteins, compared to ultracentrifugation. Finally, we applied this strategy to plasma samples of healthy individuals and those with Parkinson’s disease, identifying four potential biomarkers that provide a promising guidance for clinical diagnosis.

## 1. INTRODUCTION

Extracellular vesicles (EVs) are nanoscale vesicles secreted by most cells, which are composed of phospholipid bilayer structure, including exosome, microvesicles and apoptotic body, carrying lipids, DNA, RNA, proteins, metabolites and sugar chain complements ^1–3^. EVs are commonly found in a variety of biological body fluids, such as blood, plasma, saliva, urine, and cerebrospinal fluid. Emerging research have detected and analyzed the protein components and RNA (including microRNA) in EVs, thus recognizing that EVs are widely involved in intercellular communication and promote the transmission of molecular signals in the extracellular environment, standing a critical role in pathology of many diseases, such as cancer progression, cardiovascular diseases, and nervous system diseases ^4, 5^.

Neurodegenerative diseases are characterized by irreversible degeneration and damage of neurons in various regions of the central nervous system ^6^. Parkinson’s disease (PD) is the second most common neurodegenerative disease, affecting millions of people around the world, while its incidence increases significantly with age ^7, 8^. PD manifests with resting tremors, bradykinesia, stiffness, and postural instability ^9–11^. The main pathological manifestations of Parkinson’s disease are the decrease of pigmentation in the dense layer of substantia nigra caused by the death of dopaminergic neurons and the accumulation of α-synaptophysin in neuronal inclusion bodies ^12, 13^. Currently, treatments primarily aim to improve symptoms and enhance patients’ quality of life, while they cannot fully cure the disease. Clinical diagnosis of PD still relies on symptoms, with most patients exhibiting obvious dyskinesia at diagnosis. Therefore, the development of early diagnostic biomarkers of PD is of great significance for early treatment and improving the prognosis.

EVs can cross the blood-brain barrier and peripheral circulation, playing a crucial role in exchanging molecular information and promoting intercellular signal transduction. This function is significant in the pathogenesis of neurodegenerative diseases like PD, making EVs potential biomarkers for studying central nervous system functions and identifying neurodegenerative conditions ^14, 15^. Plasma-derived EVs are particularly noteworthy as biomarkers due to their application in fluid biopsies—a rapid and minimally invasive sampling method. This approach offers easy sampling and reduced trauma, highlighting its importance in disease biomarker research. However, the complexity and viscosity of plasma pose challenges to EVs separation. Despite the development of various methods for isolating and purifying EVs from plasma—such as ultracentrifugation, precipitation, chemical affinity purification and size exclusion chromatography ^16–19^, there are still some challenges in the separation of plasma EVs. For example, traditional methods can be time-consuming, have low separation purity, and result in co-separation of contaminants the pollutants are co-separated with EVs ^20,21^. Recent advancements have seen antibody-based affinity technologies improving EV enrichment efficiency. Priyanka Sharma et al. developed an immunoaffinity-based method for capturing melanoma-derived exosomes from the plasma of melanoma patients ^22^. Jiyun Lim et al successfully isolated exosomes from plasma of patients with breast cancer and lung cancer using antibody cocktail[conjugated magnetic nanowires ^23^. Similarly, Yoon-Tae Kang et al. used a microfluidic interface based on immune affinity to isolate and analyze exosomes in melanoma blood samples ^24^. However, EV immunoaffinity separation based on magnetic beads is usually combined with flow sorting and Western blotting (WB) analysis, but is not compatible with nanoparticle tracking analysis (NTA) or proteomic analysis ^25, 26^. Therefore, there is an urgent need to develop a fast and efficient method for EVs separation.

With the rapid development of biomarker research technology, high resolution liquid chromatography-tandem mass spectrometry (LC-MS/MS) has emerged as a key tool for the high-throughput and in-depth characterization of the plasma proteome ^27^. Furthermore, data-independent acquisition mass spectrometry (DIA-MS), an innovative bottom-up proteomic strategy, has gained widespread use in biomarker screening within large-scale clinical samples. Compared to data-dependent-analysis (DDA) method, the DIA strategy can capture all the fragment information of all parent ions in the sample without any deviation or omission. Due to its high stability and precision, DIA is particularly well-suited for studying proteomes in large sample sizes ^28, 29^.

In this study (**Figure 1**), we developed an antibody-based affinity magnetic beads, EVlent (**E**xtracellular **V**esicles iso**L**ated **E**fficiently, **N**aturally, and **T**otally), which we applied for the first time to separate EVs from small amount of plasma, demonstrating extensive compatibility with proteomic analysis. The method can identify more than 2,000 proteins from 100 μL of plasma, and even about 1,000 proteins in a tiny sample of plasma (5 μL). We utilized this technique to separate EVs from the plasma of both healthy individuals and those with PD. Combined with DIA-MS method, the potential biomarkers of PD were screened to provide help for the clinical diagnosis of such disease.

**Figure 1.**
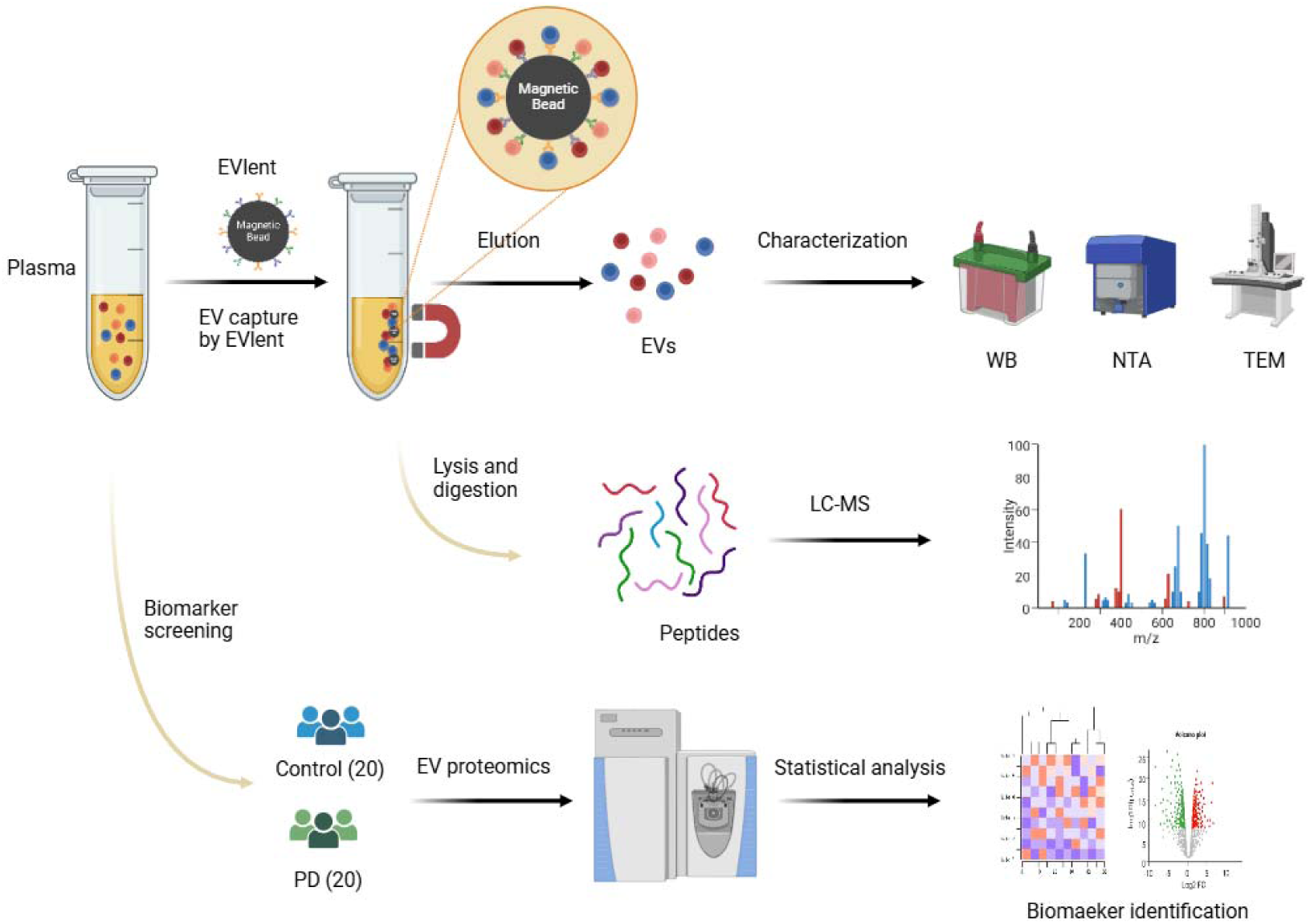
Plasma EV isolation and cargo analyses based on EVlent.

## 2. EXPERIMENTAL SECTION

### 2.1. Samples Collection

The plasma samples were collected from a tertiary-care hospital in Nanjing, Jiangsu Province. Plasma samples were obtained from healthy people and PD patients with informed consent, and all experiments were conducted in accordance with relevant guidelines and regulations. The experimental scheme has been approved (No : 2023-664-03), and the blood samples were collected using an EDTA coated vacuum cleaner. The cell fragments were initially removed by centrifugation at 4 °C (2500g for 10 mins, repeated once). After centrifugation, the plasma samples were stored at -80 °C.

### 2.2 Synthesis of affinity magnetic beads

Initially, 10 mg carboxyl magnetic beads were taken and washed three times with the coating buffer. Subsequently, the beads were combined with 0.5 mL of N-ethyl-N′-(3-(dimethylamino) propyl) carbodiimide (EDC) and 0.5 mL of N-hydroxysuccinimide (NHS) solutions, thoroughly mixed using a magnetic stirrer, and then reacted for 2 h to activate the beads. Following activation, they were again washed three times with the coating buffer. Next, appropriate amounts of anti-CD9, anti-CD63 and anti-CD81 antibodies were added to 1 mL of solution containing 10 mg resuspended magnetic beads, and mixed at room temperature for 6 h for antibody coupling. Upon completion of the coupling process, the beads were washed three times with the blocking buffer, using 1 mL of the solution for each wash. Subsequently, 1 mL of the blocking buffer was added, and the mixture was stirred at room temperature for 12 h to block remaining active sites. Finally, the blocking buffer was removed by magnetic separation, and 1 mL of magnetic bead storage buffer was added to achieve functional magnetic beads at a concentration of 10 mg/mL.

### 2.3 EVs isolation by EVlent

Take 100 μL plasma sample and dilute it 10 times with loading buffer, then add 20 μL EVlent magnetic beads, incubate the mixture at room temperature for 1 hour, and then remove the supernatant with the help of a magnetic separator. Subsequently, it was washed once with 1 mL loading buffer and twice with 1 mL PBS solution respectively. Finally, EVlent beads were eluted twice with 200 μL elution buffer (100 mM triethylamine solution (TEA)) to obtain EVs.

### 2.4 Nanoparticle Tracking Analysis (NTA)

EVs was obtained from 100 μL plasma and diluted to 1mL with PBS. Zetaview (ParticleMetrix, Meerbusch, Germany) was used to detect the particle size and concentration of separated EVs according to the standard scheme. The instrument was initially calibrated with 100nm polystyrene particles, which were diluted 250,000 times with pure water. The brightness of the instrument is set to 20, the sensitivity and shutter are set to 70 and 100 respectively.

### 2.5 Transmission Electron Microscopy (TEM)

EVs were isolated from 100 μL of plasma and resuspended in 200 μL of PBS. Subsequently, 10 μL of the solution was applied onto a 200 mesh formvar carbon-coated copper grid and allowed to dry naturally. The grid was then incubated for 3 mins with 2% phosphotungstic acid solution (pH=7.0) at room temperature for negative staining. Finally, the EVs were visualized using a Hitachi Hmur8100 electron microscope (Hitachi, Tokyo, Japan).

### 2.6 DIA-MS analysis and data processing

The peptide samples were analyzed by QE HF-X instrument (Thermo Fisher Science, USA). The liquid phase is a Thermo Science EASY-nLC 1,200 system, using a C18 chromatographic column (2.2 μm, 100 Å; Michrom Bioresources), the flow rate is 300 nL/min, the linear gradient is 53 mins, the gradient range is 8%-40% B, and the washing gradient is 7 mins (phase A: 0.1% FA, phase B: 80% ACN / 0.1% FA). The sample is scanned in DIA mode, and the parameters are as follows: Full MS resolutions:120,000, scan range: 400–1,200 m/z, AGC target: 3×10^6^, Maximum IT: 30 ms. DIA resolutions: 15,000, AGC target:1×10^6^, Maximum IT: 20ms, Isolation window: 8 m/z. Use the Direct-DIA analysis function of Spectronaut^TM^ software 18 (Biognosys, Schlieren, Switzerland) to process Thermo raw files. The analysis was conducted using the version of the human UniProt database downloaded on March 15, 2023. Use the software’s default parameters for the Direct-DIA search: set to trypsin / P, allow up to 3 missing cleavages, choose carbamoylmethylation (+ 57.02Da) as fixed modification, with methionine oxidation (+ 15.99Da) and acetylation (+ 42.01Da) as variable modification. Set the false discovery rate (FDR) for proteins and peptides to 1%. Perform quantitative analysis using MS/MS data.

### 2.7 Statistical analysis and data availability

The differential expression of the search data was analyzed by Perseus software, and the volcano map and heat map were generated (p <0.05, |log_2_(Fold change)|>1 was the differential protein). All images were drawn using GraphPadPrism (v8.0), Origin2022(v9.9) and R (4.2.3), respectively.

## 3. RESULTS AND DISCUSSION

### 3.1 Evaluation of EV isolation strategy based on affinity magnetic beads

Plasma, a commonly used fluid in biopsies, presents a significant challenge due to its complexity. Purifying or enriching EVs from plasma is crucial for biomarker discovery ^30^. CD9, CD63, and CD81 serve as key biomarkers for EVs. Despite numerous EV affinity strategies based on these markers, simultaneous modification of magnetic beads with all three antibodies remains scarcely reported. To address this, we prepared four types of affinity magnetic beads: three individually modified with anti-CD9, CD63, and CD81 antibodies, and a fourth modified with all three. To compare with the traditional ultracentrifugation (UC) separation EVs method, we first measured the efficiency of UC and four kinds of magnetic beads to separate EVs from PBS solution. The isolated EVs were analyzed by WB. As shown in **Figure 2A-2B**, where CD9 intensity was measured, the order of EV isolation efficiency obtained were CD9/CD63/CD81 (85.6%) > CD9 (81.3%) > CD81 (79.2%) > CD63 (63.6%) > UC (17.8%), indicating that anti-CD9/CD63/CD81 affinity magnetic beads overall have the highest isolation yield, while UC has the worst. Subsequently, the separation effect of affinity magnetic beads in real biological samples was evaluated. After incubating 200 μL plasma with four different magnetic beads, they were washed once with 0.01% Triton100 and twice with PBS, respectively. Then four different kinds of magnetic beads were tested by WB to observe the effect of different affinity magnetic beads capturing plasma species EVs. UC method was used as control. The analysis of **Figure 2C-2D** reveals that affinity magnetic beads modified with three antibodies demonstrate the highest effectiveness, followed by beads modified with anti-CD9, anti-CD81, and anti-CD63 antibodies individually, with the UC method being the least effectiveness. In summary, magnetic beads modified with three kinds of antibodies show superior performance, likely due to the synergistic effect of different antibody captures.

**Figure 2.**
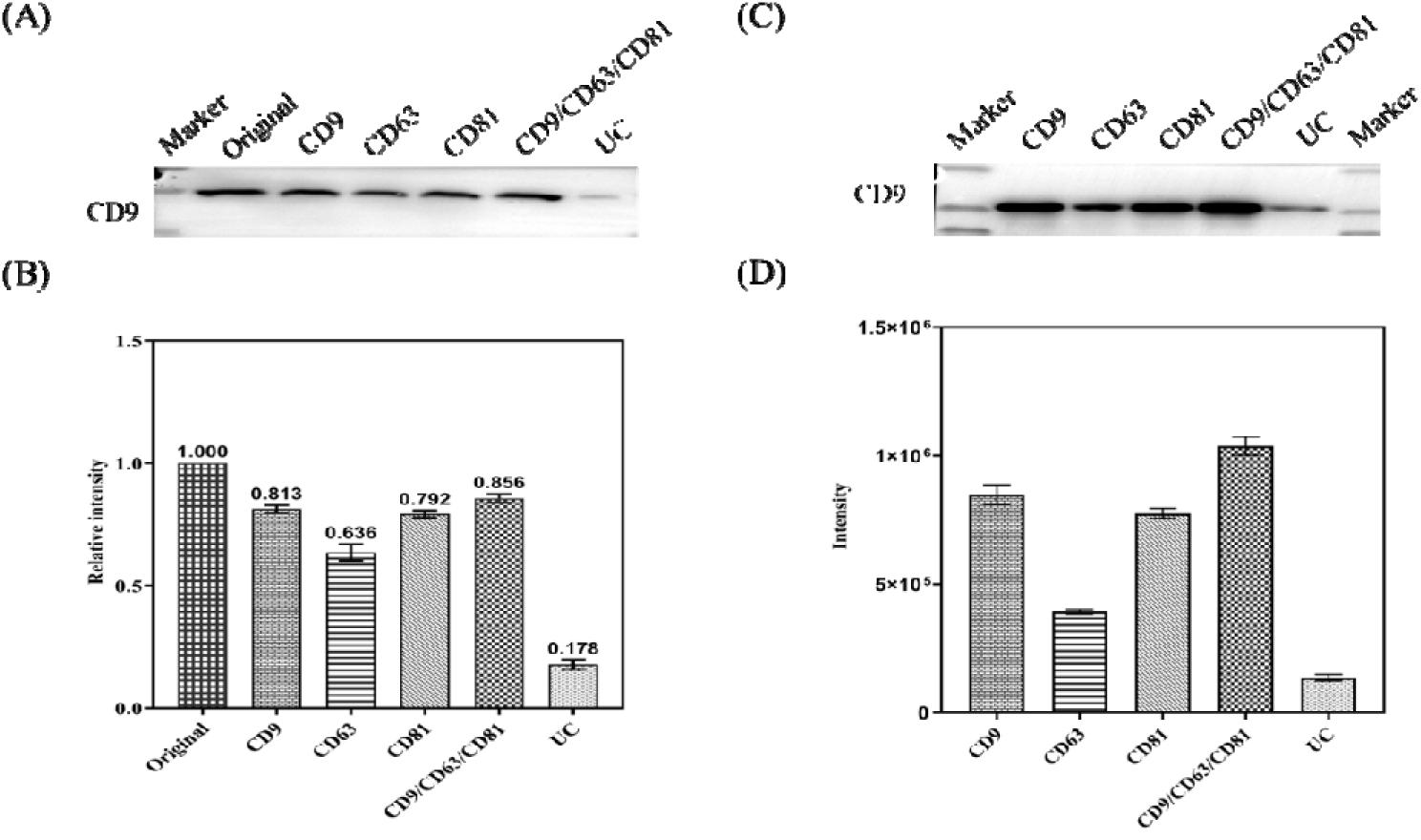
(A-B) Recovery efficiency analysis by isolating EVs in PBS and the detection of CD9 using western blotting (purified original EVs serve as the EV standard solution for recovery assay). (C) Comparison of different methods for capturing EVs in plasma; (D) Gray values of CD9 bands with different methods.

### 3.2 Full proteome comparison of captured EVs

To further evaluate the effects of different affinity magnetic beads, we compared this method with traditional UC methods. We used five plasma samples of 200 μL each, separating EVs using both affinity magnetic beads and UC, respectively. Following isolation, EVs were pre-treated and analyzed via LC-MS. In a single run, we identified 4,503 unique peptides corresponding to 449 unique proteins from the UC, 17,435 unique peptides corresponding to 2,292 unique proteins from the CD9 affinity magnetic beads, 15,437 unique peptides corresponding to 2,123 unique proteins from the CD63 affinity magnetic beads, 15,517 unique peptides corresponding to 2,122 unique proteins from the CD81 affinity magnetic beads, and finally 19,315 unique peptides representing 2,430 unique proteins from CD9/CD63/CD81 affinity magnetic beads isolation (**Figure 3A**). In addition, we compared the overlap of proteins identified by our types of affinity magnetic beads (**Figure S1A**). The figure shows that 1,772 proteins were identified by all four types of affinity magnetic beads, whereas 328 proteins were absent in the affinity magnetic beads modified with three types of antibodies. We also compared the abundance levels of three EV biomarkers (CD9, CD63, and CD81) across different affinity magnetic beads. As illustrated in **Figure S1B**, the abundance of all three EV biomarkers was highest in the affinity magnetic beads modified with three antibodies simultaneously. There is a correlation between the number of proteins identified by these affinity magnetic beads and the abundance of CD9 markers. Further comparison of the proteins detected by different methods against the top 100 proteins listed in the Vesiclepedia database showed that UC detected 42 proteins, whereas CD9/CD63/CD81 affinity magnetic beads detected up to 91 proteins (**Figure S2** and **Figure 3B**). The results demonstrate that CD9/CD63/CD81 affinity magnetic beads offer the best performance, while UC perform the worst.

**Figure 3.**
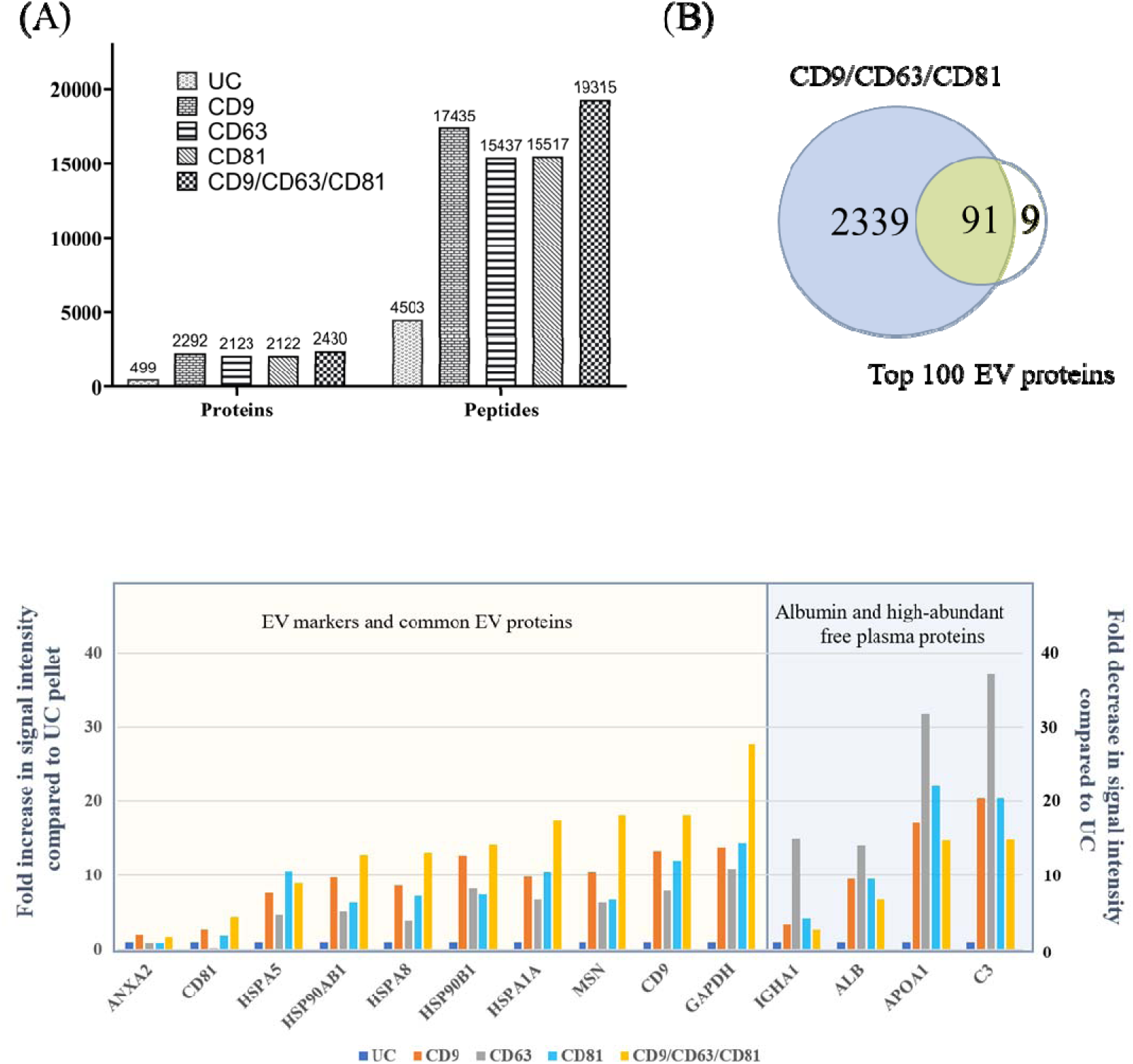
Proteomic analysis of EVs isolated by different methods. (A) Full proteome isolated by different methods; (B) Overlapping with top 100 proteins of EVs; (C) Fold increase in the total proteome intensity of 10 EV proteins (left) and fold decrease in the total proteome intensity 4 commons contaminants proteins (right) from LC-MS data compared to the UC sample.

Notably, various EVs purification techniques are susceptible to contamination by non-EV plasma constituents (such as plasma proteins, complement factors, and immunoglobulins), particularly methods reliant on chemical precipitation and precipitation ^22, 31^. Therefore, we assessed the specificity of the 4 different affinity magnetic beads by comparing the relative intensity of 10 typical EV proteins and 4 common plasma proteins in EV samples purified by the beads and UC. As shown in **Figure 3C**, samples treated with affinity magnetic beads exhibited much higher intensities of EV markers and related proteins compared to UC. Notably, in most cases, the magnetic beads modified with the three antibodies showed the highest intensities. In addition, compared with the traditional UC separation method, the intensity of four common pollutants in the affinity magnetic beads was significantly reduced in samples treated with the affinity magnetic beads. The heterogeneity of EVs from different sources in biological samples can lead to their variability in their capture ^32, 33^. When a single modified affinity magnetic bead is used for enrichment, the EVs with no or low expression of the label may be missed. Using multiple antibodies are used, the EVs in the sample can be enriched more comprehensively and the recovery rate can be improved. In summary, the affinity magnetic beads modified with three different antibodies demonstrated the best performance.

The ratio of the three antibodies on the affinity magnetic beads may influence their ability to capture EVs. To enhance the performance of these beads, we optimized the ratios of the three antibodies. Our experiments revealed that among beads modified with CD9 antibody demonstrate the best affinity for EVs, while those modified with CD63 antibody show the poorest affinity. As shown in the **Figure S3**, increasing the proportion of the CD9 antibody resulted in a higher number of identified proteins and peptides. Conversely, increasing the proportion of the CD63 antibody results in a decrease in protein and peptide identification. For the CD81 antibody, an initial increase in its proportion led to more identified proteins and peptides, followed by a decrease. This phenomenon might be due toa high concentration of antibodies increases the likelihood of non-specific binding, where antibodies may bind to non-target EVs or other plasma components. Such non-specific binding can block the active sites on the surface of magnetic beads, thereby reducing the efficiency of target EVs capture. Optimally, setting the ratio of the three antibodies (CD9:CD63:CD81) was set to 3:1:1 resulted in the highest number of proteins and peptides is identified in plasma EVs. Therefore, our focus was on the affinity magnetic beads modified with the three antibodies in this optimized ratio, which we have named EVlent (Extracellular Vesicles isolated Efficiently, Naturally, and Totally).

To further evaluate the performance of EVlent beads, we conducted a more systematic assessment. The EVlent beads’ upper surface is coated with antibodies CD9, CD63, and CD81, ensuring efficient and specific EV separation. We compared the efficacy of EVlent in capturing plasma EVs to that of UC. WB was used to detect three EV markers: CD9, TSG101, and HSP70. **Figure 4A** shows clear bands for CD9, TSG101, and HSP70 in the EVlent lane, while these bands appear weaker in the UC lane. Additionally, no Calnexin bands were detected in either separation strategy. Next, we used nanoparticle tracking analysis (NTA) to analyze the separated EVs. As depicted in **Figure 4B**, both methods yielded EVs predominantly in the 0-400 nm range. The concentrations of plasma EV obtained by EVlent and centrifugation were 2.53×10^9^ and 1.93×10^9^ particles/mL, respectively, indicating EVlent’s superior separation efficiency over UC. Further validation of successful plasma EV separation was achieved through transmission electron microscopy (TEM). The TEM images (**Figures 4C and 4D**) clearly display the intact EVs. These findings confirm the successful isolation of EVs from plasma using EVlent magnetic beads, demonstrating significantly enhanced separation efficiency compared to UC.

**Figure 4.**
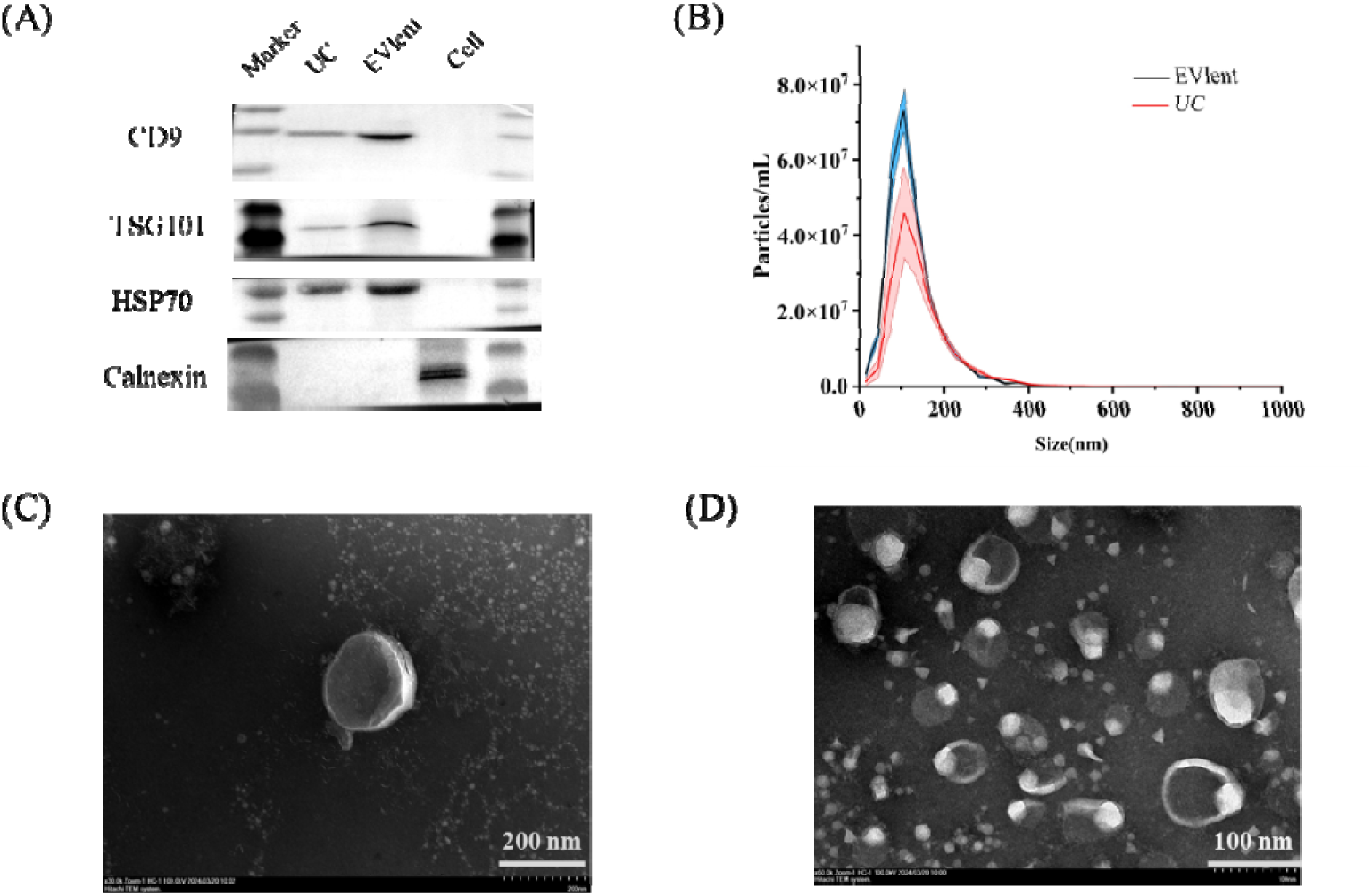
Characterization of EVs isolated from plasma by EVlent. (A) Western blot results of plasma EVs isolated by UC and EVlent. The cell extract was used as the control. CD9, CD81 and HSP70 were selected as biomarkers of EVs, and calnexin was used as a negative control; (B) NTA results of plasma EVs isolated by UC and EVlent; and (C, D) TEM images of plasma EVs from isolated by EVlent.

### 3.3 Evaluation of EVlent beads under different conditions

To investigate the optimal utilization of EVlent magnetic beads for EVs enrichment, we introduced varying volumes of these beads into 100 μL plasma samples, followed by WB analysis after EVs enrichment. As depicted in **Figure S4A**, the presence of CD9 bands remained relatively unchanged when the bead volume was reduced from 50 μL to 20 μL. However, a marked decrease in CD9 band intensity was observed with 10 μL and 5 μL bead volumes. Thus, 20 μL of EVlent magnetic beads is sufficient for complete EVs capture in 100 μL of plasma. Next, to determine the optimal incubation time for EVlent magnetic beads, six 100 μL plasma samples were incubated with 20 μL of beads for varying time (15 mins, 30 mins, 45 mins, 60 mins, 75 min, 90 mins). Following incubation, the magnetic beads were washed and EVs were obtained. The obtained EV was detected by WB analysis to observe changes in CD9 bands. As shown in **Figure S4B,** the intensity of CD9 bands gradually increases with incubation time, and when the incubation time reaches 60 mins, no obvious change in CD9 bands was observed. Therefore, the ideal incubation time of EVlent beads is 60 mins. Material stability is crucial for experimental outcomes. To facilitate high-throughput analysis of clinical samples, a method must exhibit low experimental variability. Hence, we assessed the reproducibility of the EVlent method across plasma samples. We took six identical plasma samples and enriched EVs, verified by WB experiment. The CD9 band intensity remained consistent across six replicate experiments, as illustrated in **Figure S4C**, demonstrating the robust stability of EVlent magnetic beads in plasma EVs enrichment processes. To assess plasma usage differences, we analyzed seven plasma samples with varying volumes (5 μL, 10 μL, 20 μL, 50 μL, 100 μL, 150 μL, 200 μL) using EVlent for EVs separation.

Following separation, EVs detection was performed via WB to observe CD9 band effects under varied conditions. **Figure S4D** illustrates that CD9 band intensity progressively increases with plasma volume, and a fuzzy band can be seen even in 5uL. These findings indicate that EVlent magnetic beads effectively capture EVs in plasma, even in trace samples.

### 3.4 Proteomic comparison of different plasma volumes

Plasma-based fluid biopsy is a minimally invasive technique for detecting biological markers. Plasma is a precious biological sample. Affinity-based immune-separation provides unique advantages for recovering EVs from complex and viscous liquids, including enhanced efficiency and specificity of EVs capture, integrity of isolated EVs, and selective sources ^31^. To assess the impact of different plasma volumes on protein identification, we utilized various plasma sample volumes (5 μL, 10 μL, 20 μL, 50 μL, 100 μL, 150 μL, 200 μL) to capture EVs using EVlent magnetic beads. The captured EVs were processed with phase transfer surfactant (PTS) buffer for denaturation, followed by reduction, alkylation, and trypsin digestion to produce peptides for LC-MS analysis. As shown in **Figure S5**, the number of identified proteins and peptides progressively increased with the volume of plasma analyzed. Specifically, when the plasma volume ranges from 5 μL to 100 μL, the increase is significant. However, between 100 μL and 200 μL, the rate of increase becomes more moderate. Notably, utilizing 100 μL of plasma facilitates the identification of over 2,000 proteins and 15,000 peptides. With 200 μL of plasma, we can identify as many as 2,402 proteins and 17,532 peptides. Remarkably, even with as little as 5 μL of plasma, over 800 proteins and 5,000 peptides can be identified. Compared with other reported affinity strategies, the immunoaffinity magnetic beads we developed can identify more proteins and peptides in plasma EVs ^21, 34^. These results demonstrate that EVlent has a strong capability to capture EVs in plasma without affecting downstream proteomic analysis. Given the precious nature of plasma samples, we use 100 μL plasma for follow-up proteomic studies.

### 3.5 LC-MS analysis of the EV proteome of clinical plasma samples

PD, characterized by its rapidly increasing prevalence among individuals over the age of 60, affects approximately 1% of the global population. This neurodegenerative disorder currently lacks biomarkers for early diagnosis and remains incurable. Aside from genetic detection of specific cases, diagnosis is only confirmed through the hallmark neuropathological changes in the brain post-mortem ^35, 36^. So far, there are no reliable biomarkers for clinical practice. Mass spectrometry (MS)-based proteomics emerges as a potent technique for identifying differential protein abundance levels between patients and healthy individuals, offering potential avenues for research and diagnosis ^12^. In this study, LC-MS was utilized to conduct a proteomic analysis on plasma samples from two distinct cohorts: a control group (C, n=20) and a PD patients group (PD, n=20), aiming to identify novel biomarkers for PD diagnosis. A total of 40 plasma samples, each of 100 µL, were processed. EVs were isolated using EVlent magnetic beads, followed by protein extraction in PTS buffer. Subsequent digestion with Lys-C and trypsin, detergent removal, and desalting were performed before freeze-drying the samples for LC-MS analysis. Across the cohorts, 3,921 proteins and 40,413 peptides were identified through label-free quantitative analysis.

Statistical results were obtained through in-depth data analysis, resulting in the generation of visual hierarchical clustering groups (heatmaps) and volcano maps (**Figure 5**). Hierarchical cluster analysis of quantitative proteomics was performed for all individual biological repeats (p <0.05). To visualize the quantitative results more comprehensively, volcano maps, based on t-test statistics, were used to compare and analyze the EVs proteome between every two sample categories among the three sample categories (p < 0.05, |log_2_(Fold Change)| > 1 indicated differential proteins). Compared to the control group, we identified 88 significantly up-regulated proteins and 148 significantly down-regulated proteins in PD group samples. Among these up-regulated total proteins, a number of proteins such as NDUFS4, KNG1, APOE, MMP14, TIMP2, MFGE8, MAP2K1 and so on have previously been reported to be associated with PD ^37–43^.

**Figure 5.**
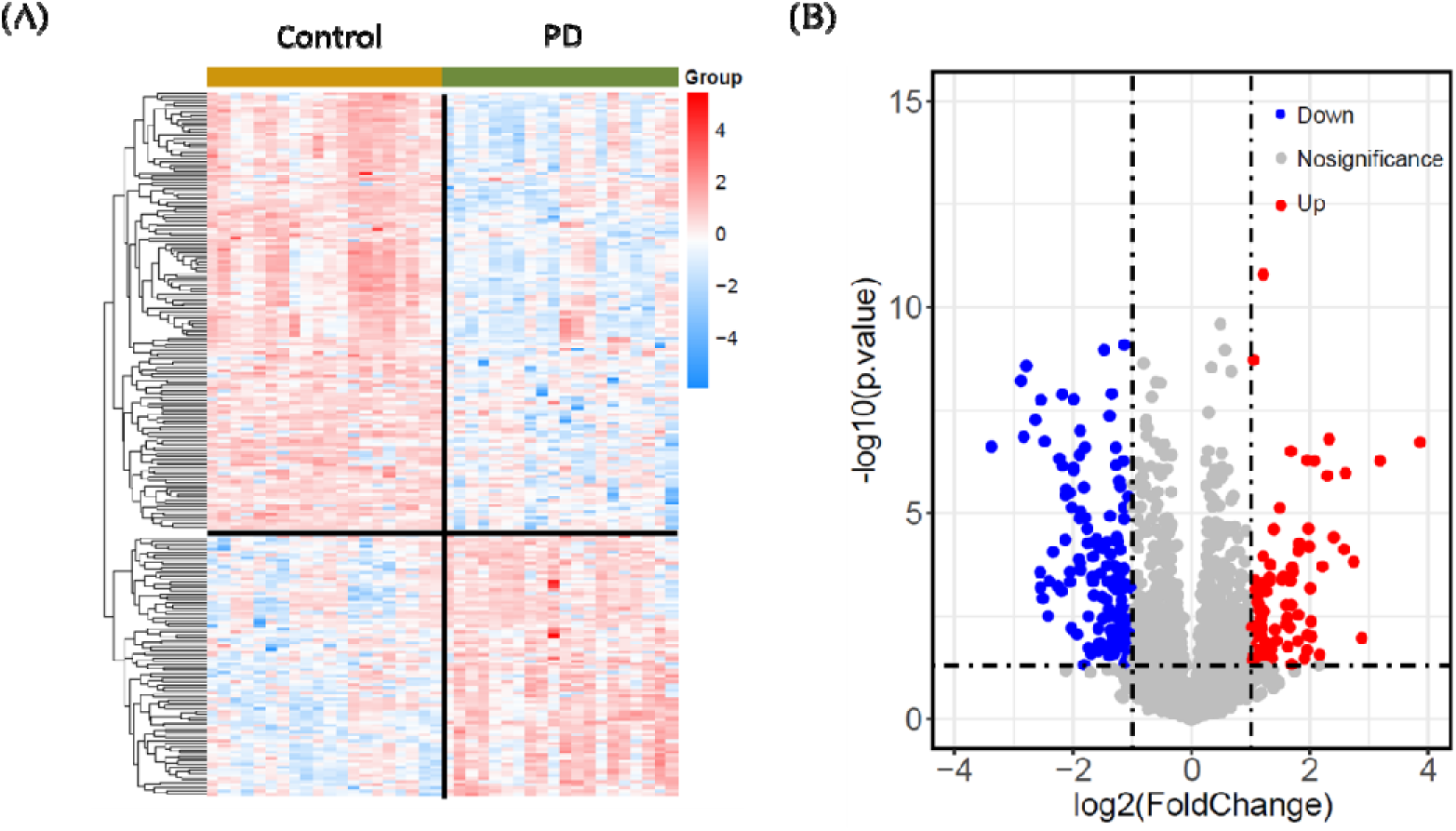
(A) Hierarchical clustering analyses on quantitative proteomics; (B) Volcano plots representing the quantitative comparison of the plasma EV proteomes (Control versus PD in the full proteome).

**Figure 6.**
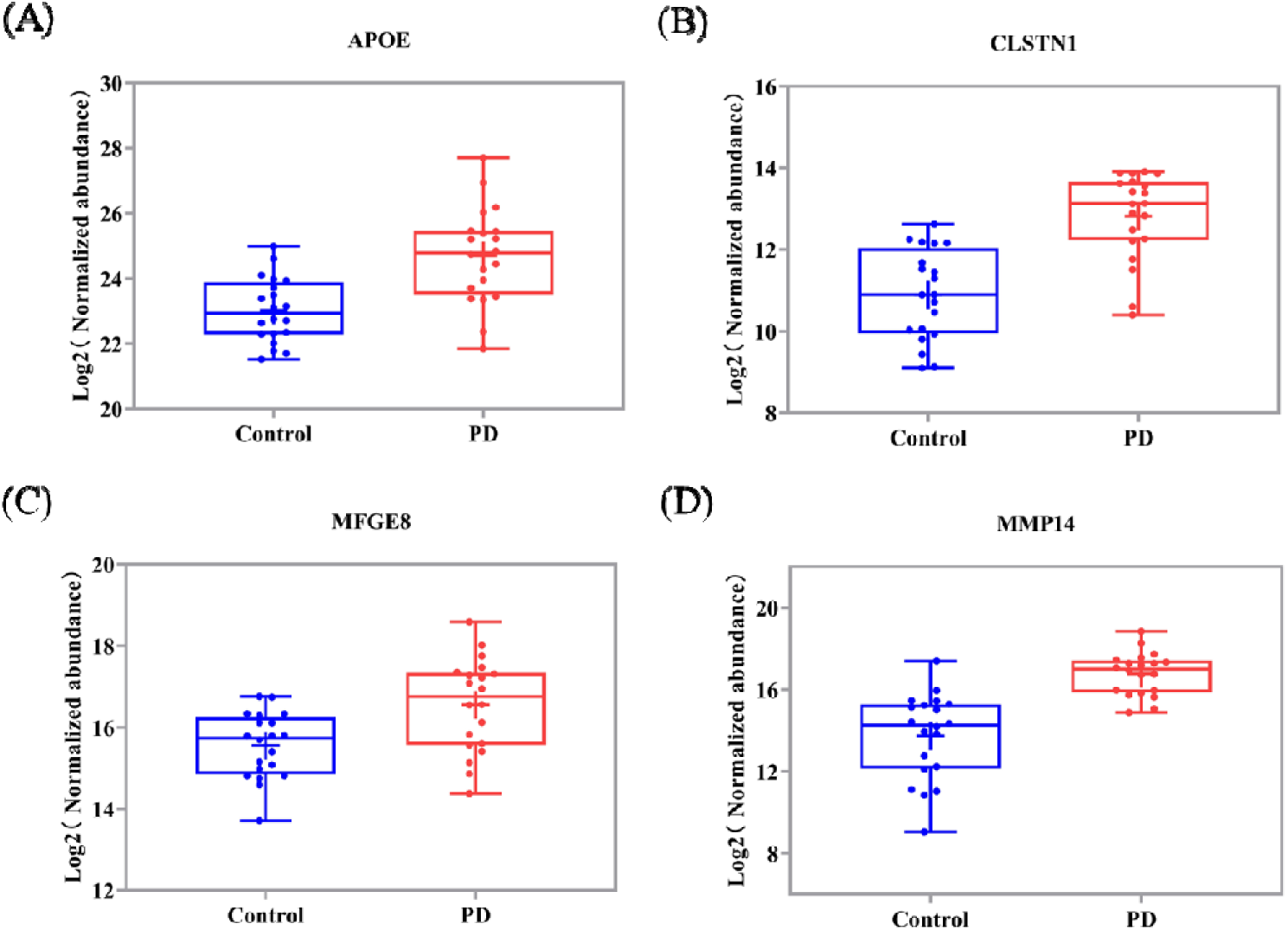
Relative abundance data of the selected four proteins as potential markers to differentiate the relevant categories. (A) APOE; (B) CLSTN1; (C) MFGE8; and (D) MMP14. All values are log_2_ conversions of protein-normalized abundance levels, as determined by LC-MS.

We focused on several proteins with significant changes, and identified the four most promising biomarker representatives. The relative abundance of these proteins, outlined in DIA model, is illustrated in the linear box diagram across the three categories. Previous studies have confirmed that these targets are associated with neurodegenerative diseases. APOE, a classical lipid-binding protein, mediates lipid transport within the central nervous system and peripheral tissues. Increasing evidence suggests that polymorphisms in the APOE gene are closely associated with Alzheimer’s disease, vascular atherosclerosis, and the regulation of human lifespan. APOE is regarded as a key susceptibility gene for age-related degenerative diseases ^44^. It has been reported that genetic variations at the APOE locus are linked to cognitive decline in PD and dementia. Furthermore, the APOE genotype may predict the cognitive trajectory in PD. ^43, 45, 46^. CLSTN1, a type I transmembrane protein and member of the cadherin superfamily of cell adhesion molecules, plays a crucial role in the nervous system. Previous studies have shown that CLSTN1 is highly expressed in cerebrospinal fluid patients with PD compared with controls ^47^. MFGE8, the primary component of milk fat globules, is secreted from the apical surface of mammary epithelial cells during lactation. It is widely expressed and functions as a secretory protein mediating various cellular interactions, such as the phagocytosis of apoptotic lymphocytes and other cells, sperm-egg adhesion, mammary gland branching morphogenesis, and enhanced tumorigenicity and cancer metastasis ^48^. Similarly, research by Kiyoka Kinugawa et al have shown that the expression of glial-derived MFGE8 is increased in the brain of patients with PD ^41^. It has been reported that Aβ induces microglia to phagocytose other viable neurons through MFGE8, leading to neuronal death. The removal of MFGE8 has been found to protect these neurons ^49^. Additionally, Ana M. Rodríguez et al. found that during neuroinflammation induced by Bacillus abortus, activated microglia can phagocytize living neurons via the phosphatidylserine-MFGE8-vitronectin receptor pathway ^50^. MMP-14, identified as the first membrane-bound matrix metalloproteinase, plays a pivotal role in the proliferation, invasion, migration, and angiogenesis of glioma cells. Recent studies have reported that MMP-14 is highly expressed in areas of the brain that show amyloid lesions and neuroinflammation ^51^. Li et al used mouse models and human samples to determine that MMP-14 overexpression is associated with familial amyloidosis polyneuropathy and progression ^42^.Taken together, these four proteins are promising as potential biomarkers for the diagnosis of PD.

## 4. CONCLUSION

In this study, we introduce a novel immune affinity-based method for isolating EVs from plasma samples. The method facilitates the rapid and efficient separation of EVs from 100 μL of plasma, enabling the identification of up to 2,000 proteins through LC-MS analysis. Even with a small amount of plasma (5μL), nearly 1,000 proteins can be identified. Moreover, this method surpasses commonly used UC methods, significantly reducing contamination from albumin and apolipoprotein in plasma. Furthermore, we used EVlent magnetic beads to capture EVs in plasma from healthy people and patients with PD, successfully identified four potential biomarkers. Overall, the strategy demonstrates excellent performance, high sensitivity and specificity in EVs separation. It shows great potential for broad applications in the early screening and diagnosis of various cancers.

## Abbreviations

EVs: Extracellular Vesicle
MS: Mass Spectrometry
WB: Western blotting
TEM: Transmission Electron Microscopy
NTA: Nanoparticle Tracking Analysis
LC-MS/MS: Liquid Chromatograph Mass Spectrometer/Mass Spectrometer
LFQ: Label Free Quantitation
PBS: Phosphate Buffer Solution
TEA: Triethylamine
FDR: False Discovery Rate
DDA: data-dependent acquisition
DIA: data-independent acquisition
PD: Parkinson’s disease.

## Author Contributions

G.Z., H.Z and Y.Z. participated in the planning, data generation and data interpretation. E.Y. contributed to the data analysis. C.M, Y.C., Y.L. and D.W. were involved in the clinical sample collection and data interpretation. Y.W synthesized antibody affinity magnetic beads. Y.Z. and G.Z. contributed in writing the manuscript. Z. X. and E.Y. was involved in funding acquisition. All authors have read and approved the manuscript.

## Acknowledgements

This project has been funded in part by the National Key Research and Development Program of China (no. 2017YFA0700404), Fundamental Research Funds for the Central Universities of China (2242022R10062/3225002202A1), Medical Foundation of Southeast University (4060692202/021), Zhishan Young Scholar Award at the Southeast University (2242023R40031). The funders had no role in the study design, data collection and analysis, decision to publish, or preparation of the manuscript.

