## Supporting Information A for "High-Efficiency Capture and Proteomic Analysis of Plasma-Derived Extracellular Vesicles through Affinity Purification"

**Support Information**

**EXPERIMENT SECTION**

**1. Materials and reagents**

Triethylamine, sodium deoxycholate (SDC), sodium lauroyl sarcosinate, (SLS), 2-Chloroacetamide (CAA), triethylammonium bicarbonate buffer (TEAB), ethyl acetate (EA), trypsin, formic acid (FA), tris (2-carboxyethyl) phosphine (TCEP) were purchased from sigma. Trypsin and endopeptidase Lys-C were purchased from EVLIXIR. Acetonitrile (ACN, Fisher Scientific), CD9 primary antibody (13403S, CST), HSP70 primary antibody (ab2787, Abcam), TSG101 primary antibody (ab125011, Abcam), Calnexin primary antibody (ab133615, Abcam), polyvinylidene difluoride (PVDF) membrane (Millipore). Bovine serum albumin (BSA), Glycine, Sodium dodecyl sulfate (SDS), 30% acrylamide-bisacrylamide, Tris Buffered Saline with Tween20 (TBST) were purchased from Biosharp. Other reagents are common laboratory reagents without specific supplier requirements.

**2. Ultracentrifugation for EV Isolation**

The frozen plasma samples were thawed in a water bath at 37 ℃, diluted 10 times with precooled PBS (4 ℃), and then centrifuged at 10,000 g for 1 h. Subsequently, the supernatant underwent further centrifugation at 100,000 g for 70 mins at 4 ℃. After removing the supernatant, pre-cooled PBS was added to resuspend the pellets, and the sample was centrifuged again for 70 mins at a speed of 100,000 g. Finally, the washed EV pellets were collected for further experiments.

**3. Western Blotting Analysis**

EVs extracted via WB were detected using four different antibodies, with cell extracts serving as controls. The obtained EVs were dissolved in SDS-PAGE sample buffer, incubated at 95 ℃ for 10 mins, and then separated on SDS-PAGE gel. Proteins were transferred to a PVDF membrane. Following the transfer, the membrane was blocked with TBST solution containing 1% BSA and then incubated overnight with primary antibodies anti-CD9, anti-HSP70, anti-TSG101, and anti-Calnexin at a 1:2,000 dilution. After the overnight incubation, the membrane was incubated with an HRP-conjugated secondary antibody in 1% BSA in TBST. Finally, the target band was visualized by Tanon5200 image analyzer.

**4. LC−MS Sample Preparation**

Firstly, the separated EVs were dissolved in a mixed solution containing 12 mM sodium deoxycholate, 12 mM SLS, 10 mM Tris-HCl, and 40 mM CAA at pH 8.5. Then, the mixture was then boiled in a water bath at 95°C for 10 minutes, followed by the addition of four times the volume of 50 mM triethylammonium bicarbonate. Lys-C enzyme was added at an enzyme-to-protein ratio of 1:100 (w/w), and the mixture was digested at 37 °C for 3 h. After 3 h, trypsin was added at an enzyme-to-protein ratio of 1:50 (w/w), and the mixture was incubated overnight at 37 °C. The next day, the sample was acidified with 10% trifluoroacetic acid (TFA) to reach a final concentration of 1% TFA, and an equal volume of ethyl acetate was added. The resulting solution was vortexed for 2 mins and centrifuged at 17,000 g for 3 mins. The upper organic phase was discarded, and the lower aqueous phase was collected and lyophilized using a freezing vacuum centrifuge (Labconco CentriVap). The desalting process was performed using a desalting membrane (3M Empore 2240-SDB-XC) according to the manufacturer's instructions, and the desalted solution was lyophilized and ready for subsequent mass spectrometric detection.

**SUPPLYMENTARY FIGURES**

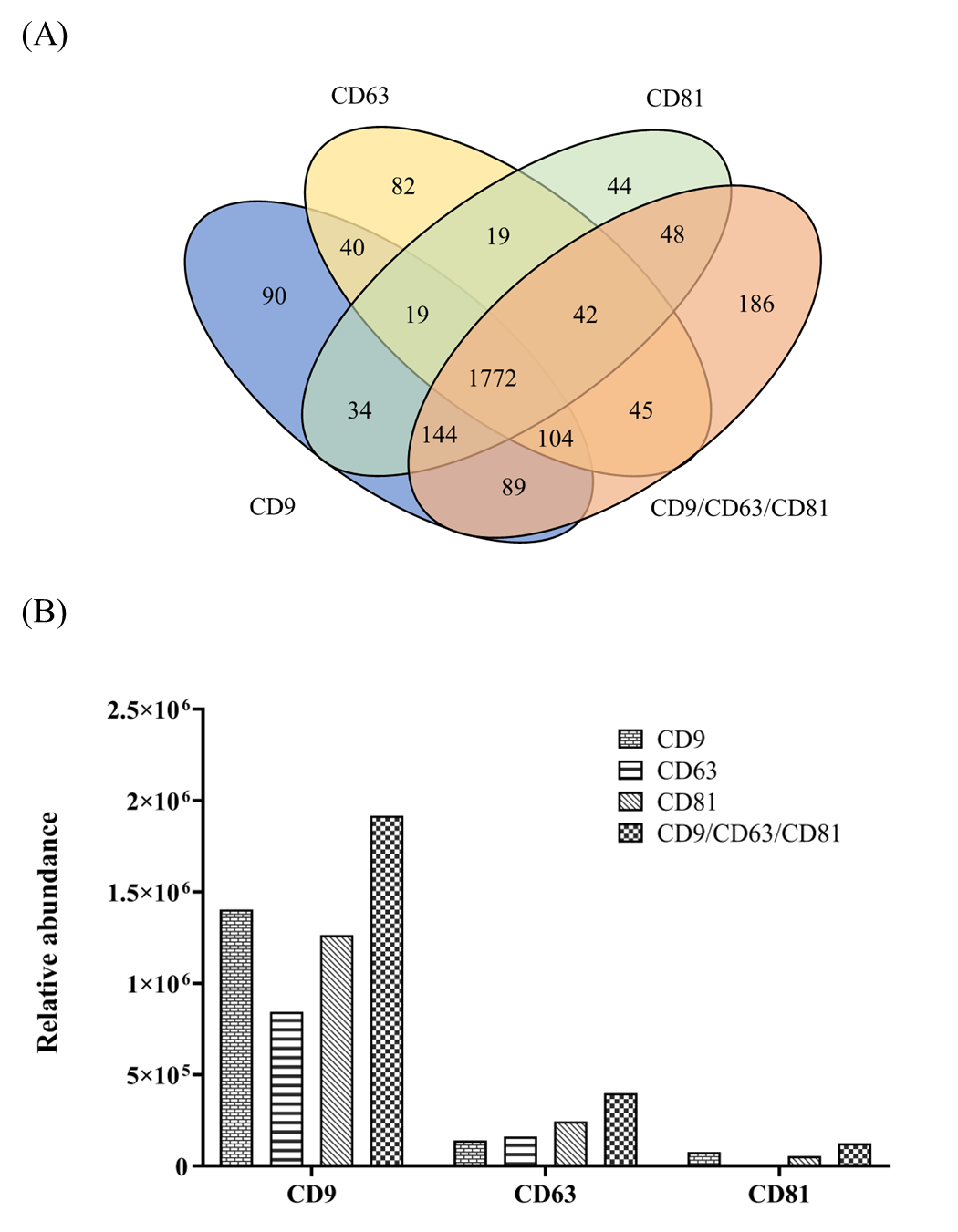

**Figure S1**. (A) Overlap of identified proteins based on LC-MS analysis from EVs isolated by 4 different affinity beads; (B) Relative abundance of 3 kinds of marker in plasma EV enriched by four kinds of affinity magnetic beads.

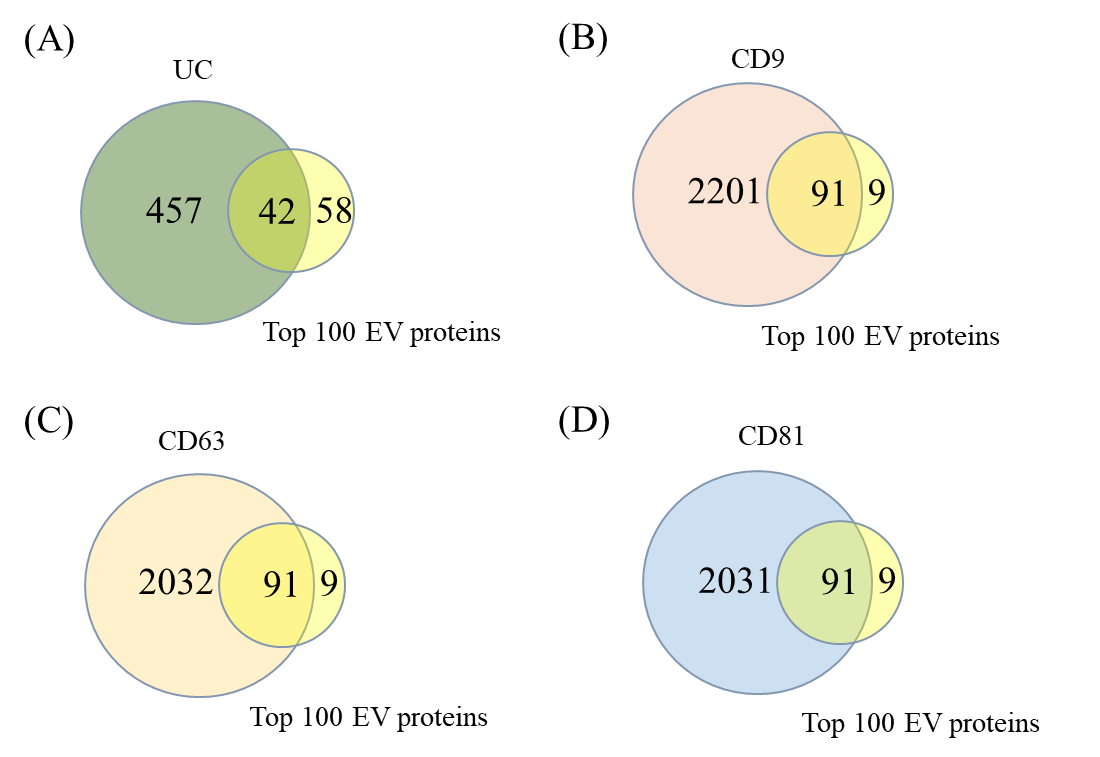

**Figure S2**. (A-D) Overlapping with top 100 proteins of EVs.

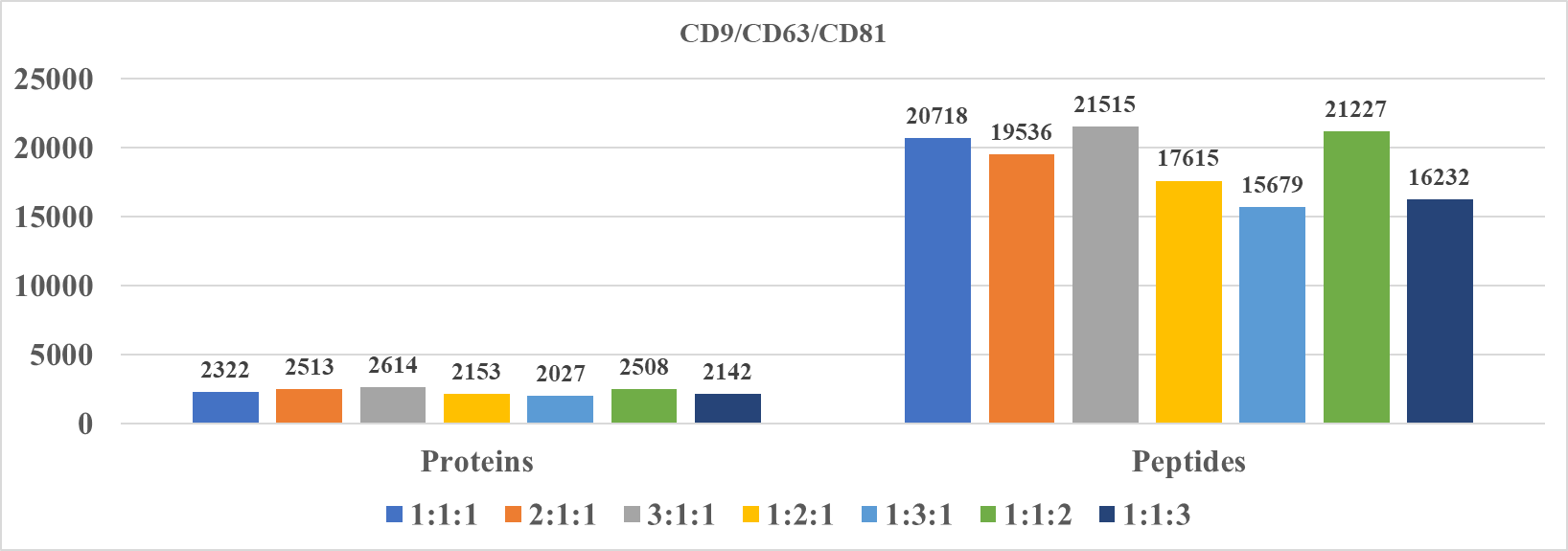

**Figure S3**. The number of proteins and peptides identified in LC-MS analysis of EVs separated by antibody modified affinity magnetic beads with different ratios.

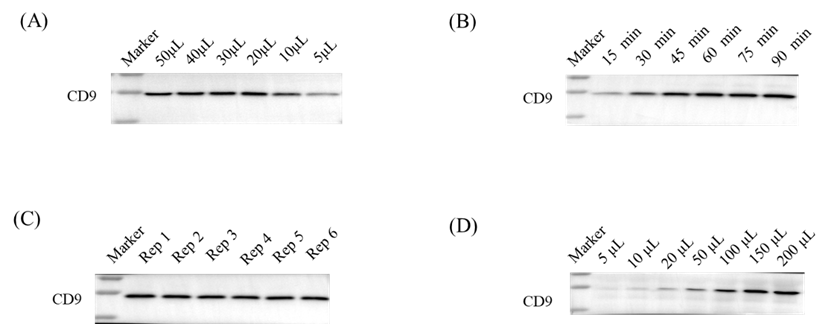

**Figure S4**. WB results of plasma EV under different conditions. (A) WB results of plasma EV separation with different volumes of EVlent; (B) WB results of EVlent beads with different incubation time; (C) Six repeated experiments on the separation of EV from plasma by EV magnetic beads; and (D) WB results of different volumes of plasma.

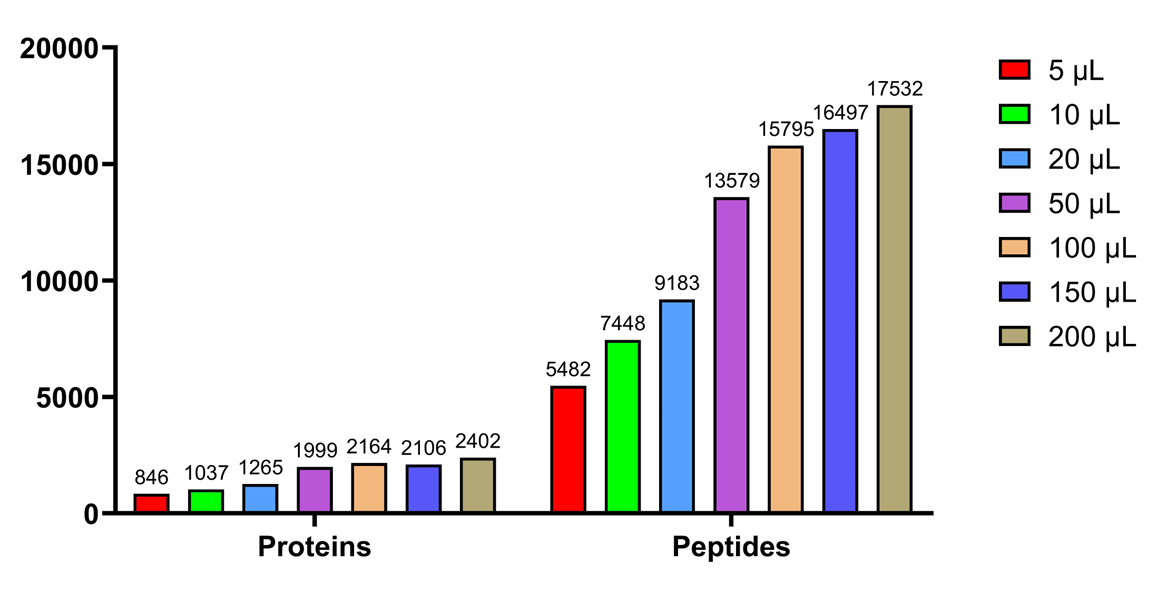

**Figure S5**. Identification of EVs in different plasma volumes by mass spectrometry.
